# Clair: Exploring the limit of using a deep neural network on pileup data for germline variant calling

**DOI:** 10.1101/865782

**Authors:** Ruibang Luo, Chak-Lim Wong, Yat-Sing Wong, Chi-Ian Tang, Chi-Man Liu, Chi-Ming Leung, Tak-Wah Lam

## Abstract

Single-molecule sequencing technologies have emerged in recent years and revolutionized structural variant calling, complex genome assembly, and epigenetic mark detection. However, the lack of a highly accurate small variant caller has limited the new technologies from being more widely used. In this study, we present Clair, the successor to Clairvoyante, a program for fast and accurate germline small variant calling, using single molecule sequencing data. For ONT data, Clair achieves the best precision, recall and speed as compared to several competing programs, including Clairvoyante, Longshot and Medaka. Through studying the missed variants and benchmarking intentionally overfitted models, we found that Clair may be approaching the limit of possible accuracy for germline small variant calling using pileup data and deep neural networks. Clair requires only a conventional CPU for variant calling and is an open source project available at https://github.com/HKU-BAL/Clair.

## Introduction

Fast and accurate variant calling is essential for both research and clinical applications of human genome sequencing^1,2^. Algorithms, best practices and benchmarking guidelines have been established for how to use Illumina sequencing to call germline small variants, including single-nucleotide polymorphisms (SNPs) and insertions/deletions (indels)^3-6^. In recent years, single-molecule sequencing (SMS) technologies have emerged for a variety of important applications^7^. These technologies, which are also known as the third-generation sequencing technologies, generate sequencing reads two to three orders of magnitude longer than Illumina reads (10–100kbp versus 100–250bp). The long read length has made the new SMS technologies, including Pacific Biosciences (PacBio) and Oxford Nanopore Technology (ONT), unprecedentedly powerful for resolving complex genome assembly problems and for detecting large structural variants^8^. However, currently available SMS technologies also have a significantly higher base error rate of 3–15%^9^, making the variant calling methods previously designed for Illumina sequencing inapplicable to SMS technologies. The lack of accurate tools for efficient variant calling has limited SMS technologies from being applied to the many problems that require SNPs and small indels.

In our previous work, we developed Clairvoyante^10^, a germline small variant caller for single molecule sequencing data. Clairvoyante does not require sequence assembly and calls variants directly from read alignments. Clairvoyante adopts a deep convolutional neural network, so that by using the truth variants called and orthogonally verified in seven human individuals by the Genome In A Bottle (GIAB) consortium^11-13^, Clairvoyante can be trained for variant calling on any new type of sequencing data without the need to look into its error profile and build a hand-crafted model. Clairvoyante takes pileup data as input and runs quickly. However, Clairvoyante’s design is unable to call multiallelic variants or indels longer than four bases. These defects remain to be solved. Meanwhile, the limit of using pileup data and deep neural networks for variant calling remains to be explored.

In this study, we present Clair, a fast and accurate system for germline small variant calling using single molecule sequencing data. With an entirely different network architecture and learning tasks (i.e. output components), Clair resolves the multiallelic and long indel variant calling problems that have prevented Clairvoyante from calling all types of small variants. We describe in detail the methods we tried that either worked or did not work for improving Clair’s performance. For ONT datasets^14^, our experiments on whole-genome variant calling in GIAB samples show that Clair outperforms Clairvoyante and other variant callers, including Longshot^15^ and Medaka^16^, in terms of precision, recall and speed. For high accuracy reads, including both PacBio CCS (Circular Consensus Sequencing)^17^ and Illumina datasets^13^, DeepVariant^18^ had modestly improved F1-scores over Clair by .11% to .13%, although Clair was seven times faster. Looking into the false positive (FP) and false negative (FN) variants of the three sequencing technologies showed that except for variants with insufficient coverage by chance, most of the others could be resolved using complete read alignments instead of pileup data or else could not be resolved at all, even with a manual inspection.

## Results

### Overview of Clair

Clair is a four-task, five-layer recurrent neural network with two bi-directional LSTM layers followed by three feedforward layers (**Figure 1**). Clair takes a BAM file as input to find candidate variants with any minor allele frequencies larger than a threshold (typically between 0.1 and 0.2), and then computes a pileup of the candidates and converts the summaries into a tensor. In a tensor, the allelic counts of bases and gaps on both strands of a candidate variant and its 16 flanking bases are encoded into 1,056 integer values. More details and pseudo code are available in the Methods section. As discussed in the Clairvoyante paper, one major unsolved problem was how to support the calling of multi-allelic variants (i.e., variants with two alternative alleles). In Clair, the problem is solved by using four new (deep learning) tasks that are entirely different from Clairvoyante. These are: 1) a 21-genotype probabilistic model with 21 probability outputs; 2) the use of three probabilities for the input, including a homozygous reference (0/0 genotype), a heterozygous variant (0/1) or a homozygous variant (1/1); 3) the length of the first indel allele, with 33 probabilities representing a length of ‘<-15bp’, ‘-15bp’, ‘-14bp’, …, ‘-1bp’, ‘0bp’, ‘1bp’, …, ‘15bp’, ‘>15bp’; and 4) the length of the second indel allele. The 21-genotype probabilistic model can represent all possible genotypes of a diploid sample at the genome position. The length of indels longer than 15bp cannot be directly inferred from the third and fourth tasks, so Clair includes an additional step that re-scans the alignments. More details on each of these steps can be found in the Methods section. The four tasks make their own decisions and are designed to cross-validate each other. For example, task two is a coarse-grained version of task one and can veto the decision made by task one. Tasks three and four should indicate 0bp indel length if an SNP variant is decided by task one. More details on how the four tasks make a joint decision are available in the Methods section. We used the ‘focal loss’ deep-learning technique to solve the problem of unbalanced variant types in training data. We used the ‘cyclical learning rate’ deep learning technique to achieve the maximum possible variant calling performance and speed up the training process to be able to handle larger training datasets. To improve Clair’s performance at lower sequencing coverages, we augmented the training data with 10 subsampled coverages of each dataset. The parameters of these three new techniques are in the Methods section.

**Figure 1.**
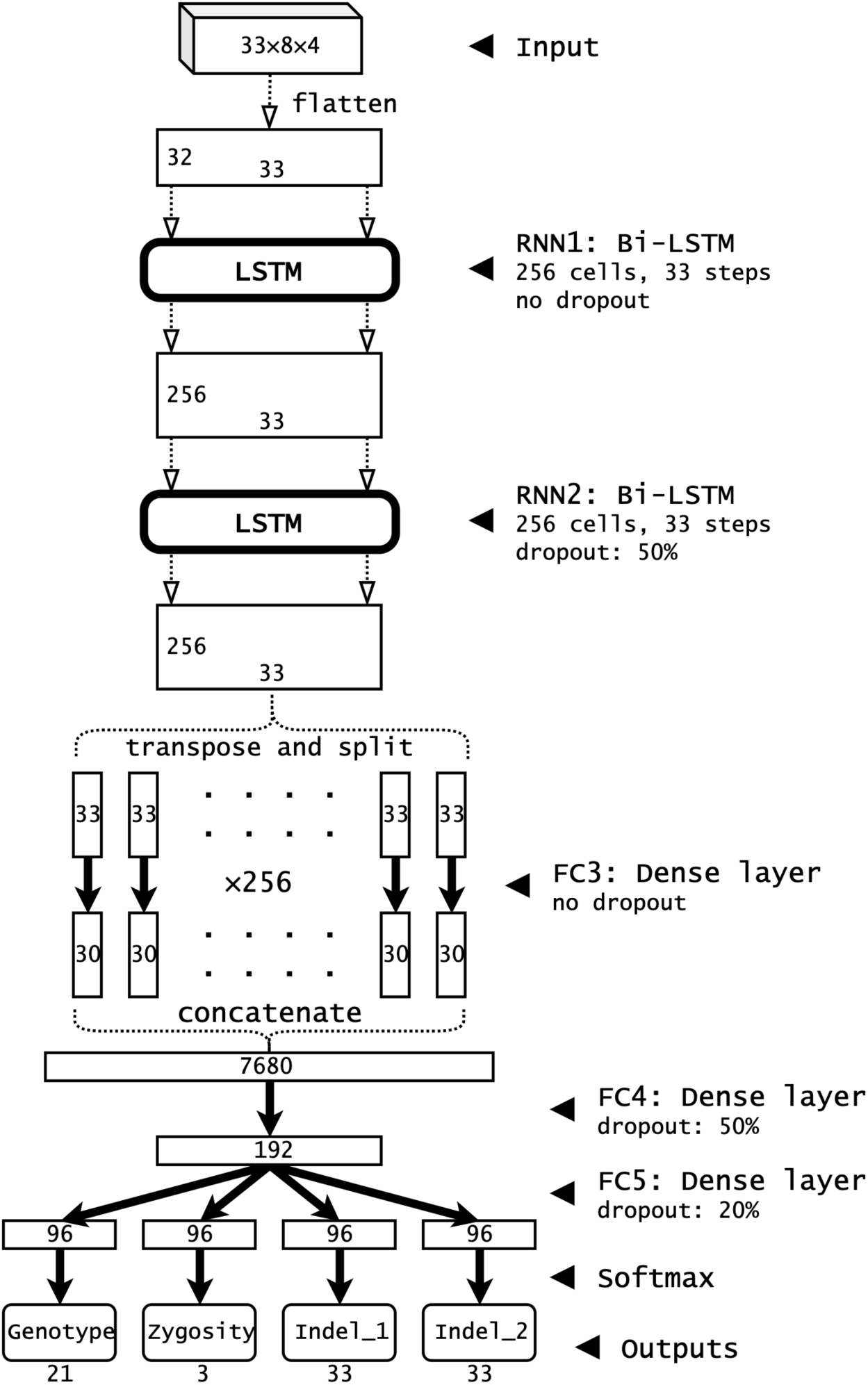
Clair network architecture and layer details. RNN: Recurrent Neural Network. FC: Fully Connected layer. Bi-LSTM: Bi-directional Long Short-Term Memory layer.

Clair has 2,377,818 parameters, which is 45.7% more than Clairvoyante (1,631,496 parameters) but only one tenth as many as DeepVariant (23,885,392 parameters). In terms of variant calling speed, Clair takes about 30 minutes, 1.5 hour, and 5 hours for a 50-fold coverage WGS sample using Illumina, PacBio CCS and ONT data, respectively, using 24 CPU cores. In our experiments, Clair was 10–20% slower than Clairvoyante, but significantly faster than DeepVariant, Longshot and Medaka.

The Methods section includes a description of procedures to augment the training data or improve Clair’s network architecture that we tested but that did not improve precision and recall of variant calling. Developers working on further improving Clair’s performance can save time by avoiding the same methods, or the same settings in a method.

### Performance on ONT

ONT datasets are currently available for two GIAB samples, HG001 and HG002. The HG001 rel6 dataset generated by the Nanopore WGS Consortium^14^ contains approximately 44.3-fold coverage of human genome (the dataset is also referred to as 1:44x, where ‘1’ means the sample suffix and ‘44x’ means the coverage). The rel6 dataset was base-called with Guppy 2.3.8, using the HAC (High-ACcuracy) model. In addition to the rel6 dataset, we obtained a separate 124.1-fold coverage dataset for HG001 (1:124x) directly from Oxford Nanopore (Philipp Rescheneder, personal communication). That dataset was base-called with Guppy 2.2.3 using the Flip-Flop model. In some experiments, we combined 1:44x and 1:124x to form a new dataset 1:168x to maximize the coverage. For HG002, we used a dataset with ∼64-fold coverage (2:64x) from the GIAB consortium, which was base-called with Guppy 2.3.5 using the Flip-Flop model. The links to the datasets are available in the Supplementary Notes. The details about “the GIAB truth variant datasets”, “removing GA4GH (The Global Alliance for Genomics and Health) low-complexity regions^6^ from benchmarking”, and “the benchmarking methods and metrics” are available in “Methods – Benchmarking”.

Figure 2 shows the precision and recall of Clair and other variant callers on SNPs and indels in multiple experiments with ONT data. Supplementary Table 1 contains more details, including precision, recall and F1-score in five categories, including overall, SNP, indel, insertion, and deletion. Our results show that Clair not only outperformed other variant callers, including Clairvoyante, Longshot, and Medaka, but also ran much faster. Using 1:168x|2:64x (i.e., test variant calling using HG002 with 64-fold coverage against a model trained using HG001 with 168-fold coverage) as Clair’s primary result, Clair achieved 98.36% precision, 96.46% recall, and 97.40% F1-score overall performance. In terms of SNPs, the three metrics were 99.29%, 97.78% and 98.53%, respectively. For indels, they were somewhat lower at 81.15%, 73.88%, and 77.34%. Clair significantly outperformed its predecessor Clairvoyante on both SNP and indel calling (overall F1-score 97.40% versus 93.45%). Clair had a slightly higher F1-score on SNPs than Longshot (98.53% versus 98.41%), but Longshot detects only SNPs, and Clair ran five times faster than Longshot (320 versus 1,797 minutes). Clair had a better performance than Medaka (overall F1-score 97.40% versus 94.81%) and ran 30 times faster (320 versus 10,817 minutes). It is worth mentioning that we didn’t benchmark Nanopolish^19^, which is also capable of variant calling on ONT data, because it also requires raw signals as input, which are not publicly available for HG002.

**Figure 2.**
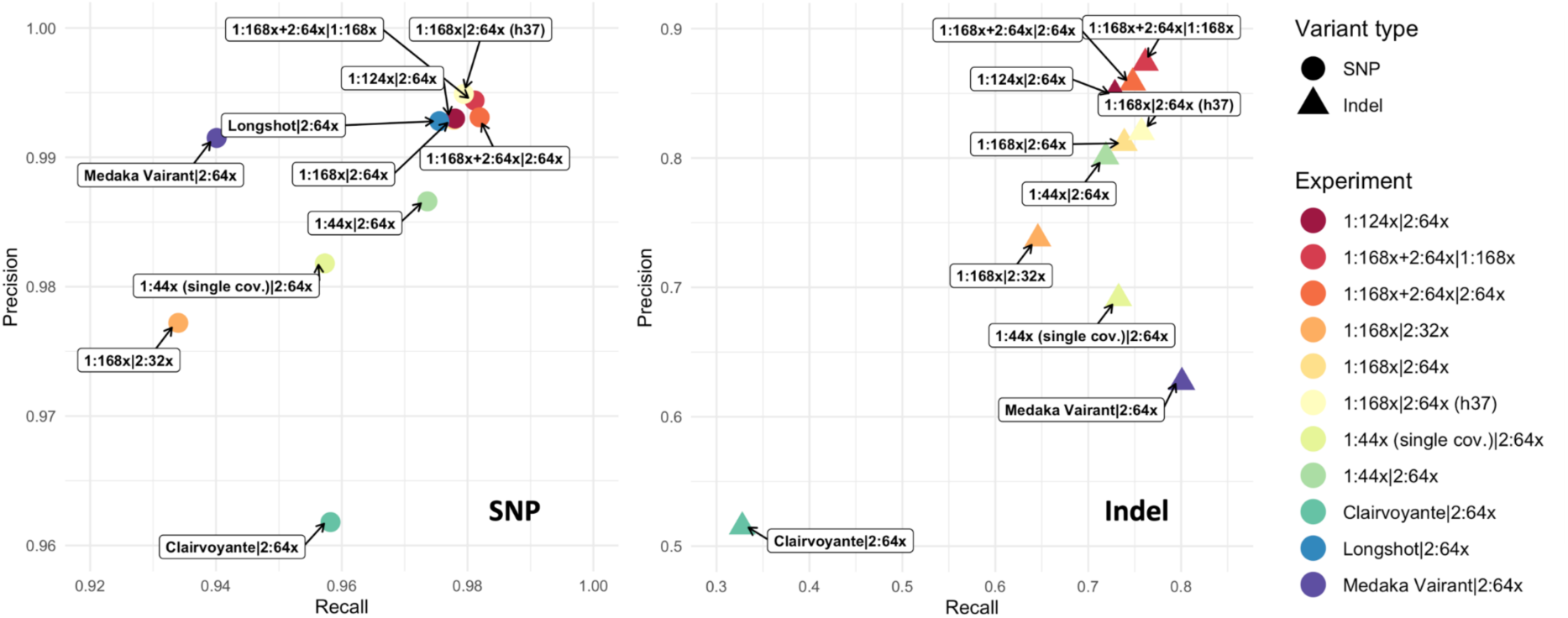
ONT benchmarking results. For Clair, the datasets used for model training and testing are separated with a vertical bar ‘|’, and are written as ‘*a:b*x’, where *a* denotes the suffix of the GIAB sample ID (e.g., 1 means HG001), and *b* denotes the coverage of the dataset. Longshot calls only SNP variants, so it is not shown in the indel results.

We ran further experiments to answer five additional questions about Clair, as follows.

#### Is the Clair model reference-genome specific?

In our experiments, performance did not depend on whether we used GRCh37 or GRCh38. The performance of 1:168x|2:64x and 1:168x|2:64x(b37) was similar; the latter experiment tested HG002 GRCh37 read alignments on a model trained using HG001 GRCh38 read alignments. Actually, 1:168x|2:64x(b37) performed slightly better than 1:168x|2:64x, with a 0.18% better F1-score on SNPs, and 1.4% on indels.

#### Does higher coverage in the test sample helps improve variant calling performance?

Yes, but improvement seems to asymptote at ∼60-fold coverage. In a comparison of 1:168x|2:64x to 1:168x|2:32x, the overall F1-score increased from 94.10% to 97.40% (+3.30%), the SNP from 95.51% to 98.53% (+3.02%), and the indel from 68.87% to 77.34% (+8.47%). Further increasing the coverage in the test sample will note significantly increase the variant calling performance as we discuss below.

#### Does higher coverage for model training help improve variant calling performance?

Yes, but it depends on the coverage of the test sample. In a comparison of 1:124x|2:64x to 1:44x|2:64x, the overall F1-score increased from 96.84% to 97.51% (+0.67%), the SNP from 98.01% to 98.54% (+0.53%), and the indel from 75.78% to 78.44% (+2.66%). In a comparison of 1:168x|2:64x to 1:124x|2:64x, the performance was similar, or even slightly dropped from 97.51% to 97.40% overall. One possible reason is that the lower coverage test sample cannot benefit from the much higher coverage used for model training. We propose how to deal with excessively high coverage in test samples (i.e., coverage exceeding that used in model training) in the Discussion section below.

#### Does multiple subsampled coverage for model training improved variant calling performance?

Yes. in a comparison of 1:44x|2:64x to ‘1:44x (single cov.)|2:64x’, the latter used only the full coverage 44-fold in model training; the overall F1-score increased from 95.47% to 96.84% (+1.37%), the SNP from 96.94% to 98.01% (+1.07%), and the indel from 75.78% to 78.44% (+2.86%). The results show that even without sufficient coverage for model training, using multiple subsampled coverage still improved the variant calling performance significantly.

#### What is the upper bound on performance?

To determine Clair’s performance cap using the current ONT data, we intentionally overfitted Clair by adding the samples we are going to test to the model training. Even though Clair is designed with multiple generalization techniques, including ‘dropout’ and ‘L2 regularization’, exposing the test samples to model training is a biased evaluation, and if a true variant is not called even after this biased training, this suggests the input signal is simply too weak. The two tests we did were 1:168x+2:64x|2:64x and 1:168x+2:64x|1:168x. Although the test sample coverage in the first test was much lower than that in the second (64-fold against 168-fold), their performance was similar, with the overall F1-score at 97.77% and 97.82%, SNP at 98.75% and 98.77%, and indel at 79.92% and 81.37%. The biased test 1:168x+2:64x|2:64x did not significantly outperform 1:168x|2:64x; the overall F1-score increased from 97.40% to 97.77% (+0.33%), SNP from 98.53% to 98.75% (+0.22%), and indel from 77.34% to 79.92% (+2.58%). Even with this biased experiment, we observed that the performance of using Clair on the current ONT data was capped at about 97.8% F1-score overall, 98.8% on SNPs, and 80% on indels. We consider how the new ONT chemistry that provides a lower base error rate can raise the upper bound of Clair’s variant calling performance in the Discussion section below.

We analyzed and categorized the FP and FN results of Clair on ONT data. We randomly extracted 100 FPs and 100 FNs from the 1:168x|2:64x experiment. **Figure 3** shows a summary and examples of different categories, and Supplementary Table 2 shows a detailed analysis of each FP and FN. Within the 100 FPs, the three largest categories are “Incorrect allele with AF≥0.2” (41/100), “Homopolymer” (25/100), and “Tandem repeat” (11/100). “Incorrect allele with AF≥0.2” means that at the FP variant, an incorrect allele dominates other alleles in the read alignments (including the correct one), and the incorrect allele has a frequency ≥20%. “Homopolymer”, “Tandem repeat”, and “Low complexity region” mean that the FP variant is in a repetitive region, which remains difficult for ONT base-calling. It is worth mentioning that these repetitive regions are ≤10bp because we removed all GA4GH low-complexity regions longer than 10bp from benchmarking. It may not be possible to perfectly resolve these three categories for FP variants using pileup data for variant calling, although complete read alignments might help to provide better precision. Three out of 100 FPs had “Incorrect insertion bases”, while two out of 100 were categorized as “Overlapping insertions”, which means that the alleles of two consecutive insertions overlapped each other in an input tensor; thus, the correct allele cannot be resolved for both insertions. These two categories of errors can be resolved using the ‘--pysam_for_all_indel’ option in Clair, but this slows down Clair for ONT data by a factor of up to ten times. Other errors, including “Incorrect indel length” and “Incorrect zygosity”, are errors made by Clair’s neural network. In the 100 FNs, the three major categories are “Correct allele with AF<0.25” (54/100), “Homopolymer” (18/100), and “Tandem repeat” (7/100). “Correct allele with AF<0.25” means that at the location of the missed (FN) variant, the signal of the correct allele is rather weak, with allele frequency lower than 25%. One FN categorized as “More than two possible alternative alleles” is an error due to an alignment error in segmental duplications, in which more than two alternative alleles seem correct.

**Figure 3.**
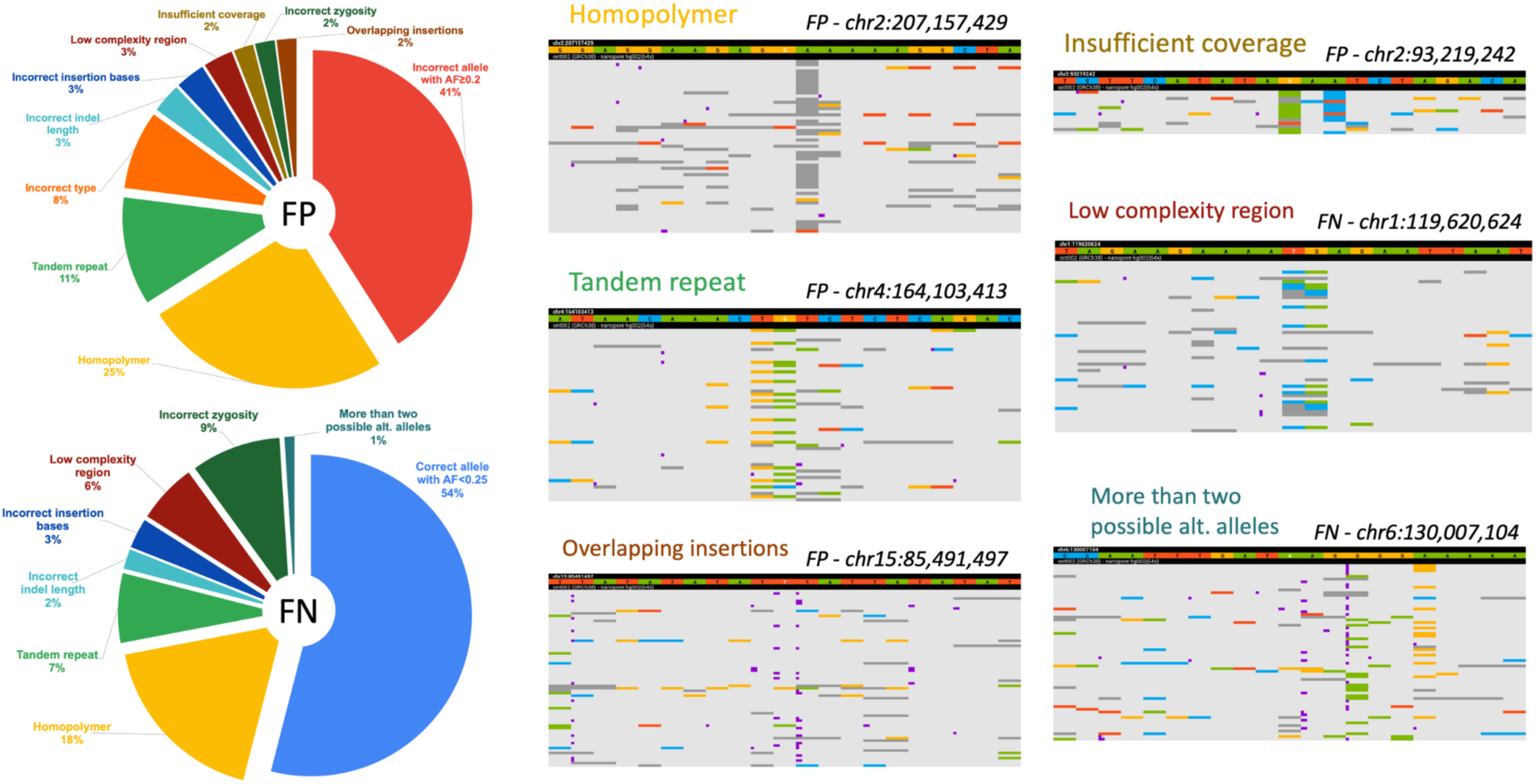
The category distribution of FPs and FNs made by Clair in the 1:168× |2:64× experiment on ONT data, and six genome browser screen captures showing examples of different categories. In the screen captures, bases A, C, G, and T are green, blue, yellow, and red, respectively. Gaps (i.e., deletions) are dark gray. Insertions are purple dots between two bases and are wider when the insertion is longer.

### Performance on PacBio CCS

In early 2019^17^, PacBio developed a protocol based on single-molecule, circular consensus sequencing (CCS) to generate highly accurate (99.8%) long reads averaging as much as 13.5kb. PacBio published CCS datasets for HG001 (in this section also referred to as 1:30x; 1 as the sample suffix and 30x means 30-fold coverage), HG002 (2:33x) and HG005 (5:33x). All three samples are involved in model training. To demonstrate a possible overfitting phenomenon on deep learning based variant callers, both HG002 and HG005 are used in benchmarking.

Supplementary Table 3 shows the results of Clair and three other variant callers: Clairvoyante, Longshot, and DeepVariant. Testing on HG002, DeepVariant performed the best, with an overall F1-score of 99.96%, SNP of 99.97%, and indel of 99.92%. The primary result of Clair 1:30x+5:33x|2:33x had an overall F1-score of 99.83%, which was 0.13% lower than DeepVariant, but outperformed both Clairvoyante and Longshot. On SNP, 1:30x+5:33x|2:33x had an F1-score of 99.88%, which was 0.09% lower than DeepVariant, 0.43% higher than Longshot, and 0.17% higher than Clairvoyante. On indel, 1:30x+5:33x|2:33x had an F1-score at 99.07%, which was 0.85% lower than DeepVariant, but 19.17% higher than Clairvoyante, showing that the new methods applied to Clair have effective solved the indel-calling problem in Clairvoyante. In terms of speed, Clair (147 minutes) is slightly faster than Longshot (206 minutes), and about seven times faster than DeepVariant (1,072 minutes). We also tested HG005. Interestingly, while Clair, Clairvoyante, and Longshot all performed better on HG005 than HG002, DeepVariant performed worse. Comparing 1:30x|2:33x to 1:30x|5:33x, Clair’s overall F1-score increased from 99.77% to 99.80%. Clairvoyante’s overall F1-score increased from 98.61% to 98.70%. Longshot’s SNP F1-score increased from 99.45% to 99.46%. The performance of the three callers verifies the quality of the HG005 dataset. However, DeepVariant’s F1-score dropped from 99.96% to 99.92%, the SNP F1-score decreased from 99.97% to 99.93%, and the indel F1-score dropped most significantly, from 99.92% to 99.78%. The most probable reason is that, DeepVariant’s current PacBio CCS model was trained completely using HG002^20^. We suggest using DeepVariant’s result on HG005 as its real performance on PacBio CCS data. The biased test 1:30x+2:33x+5:33x|2:33x found the performance cap of Clair at 99.88% on SNP, which was the same as 1:30x+5:33x|2:33x, and 99.28% on indel, which was 0.21% higher than 1:30x+5:33x|2:33x. While in 1:30x+5:33x|2:33x, the highest coverage used for model training was only 33x, we expect to fill the performance gap on indel calling by using higher coverage for model training. The performance gap between Clair and DeepVariant on HG005 (99.28% against 99.78%, −0.5%) is the result of Clair using pileup data, while DeepVariant uses complete read alignments that contain information at a per-read level. This is also a reason DeepVariant runs slower than Clair. We discuss the possibility of improving Clair to use complete read alignments without slowing down performance in the Discussion section below.

### Performance on Illumina

Approximately 300x coverage in 148-bp Illumina paired-end read data is available for five GIAB samples, including HG001, HG002, HG003, HG004 and HG005^11^. We used HG001, HG003, HG004, HG005 for model training, and HG002 for benchmarking. To resemble the typical coverage in whole genome sequencing, we used full coverage of HG001 (306-fold) and HG005 (352-fold), but down-sampled HG002, HG003 and HG004 to 52-, 57-, and 66-fold.

Supplementary Table 4 shows the results of Clair and DeepVariant. DeepVariant performed better, with an overall F1-score of 99.94%. The primary result of Clair 1:306x+3:57x+4:66x+5:352x|2:52x was an overall F1-score of 99.83%, which was 0.11% lower than DeepVariant’s. For SNPs, the F1-score of Clair was 0.09% lower than that of DeepVariant (99.85% versus 99.94%). For Indel, the F1-score of Clair was 0.42% lower than DeepVariant’s (99.48% versus 99.90%). In terms of speed, Clair was about seven times faster than DeepVariant (77 versus 537 minutes). The biased test 1:306x+2:52x+3:57x+4:66x+5:352x|2:52x found the performance cap of Clair to be 99.87% for SNPs, which was 0.02% higher than the primary result, but 0.07% lower than that of DeepVariant, and 99.57% for indels, which was 0.09% higher than the primary result, but 0.33% lower than that of DeepVariant. Similar to the ONT and PacBio CCS experiments, we expect to fill in the performance gap through partially making use of complete read alignments, as discussed in the Discussion section.

## Discussion

In this paper we present Clair, a germline small variant caller for single molecule sequencing data. The name Clair means ‘clear’ in French, echoing its predecessor, named Clairvoyante, meaning ‘clear seeing’. Clair adds new methods to solve problems that Clairvoyante had trouble with, including multiallelic variant calling and long indel calling. In our experiments on ONT data, Clair outperformed all existing tools in terms of precision, recall and speed. On PacBio CCS and Illumina data, Clair performed slightly worse than DeepVariant, but ran about an order of magnitude faster. Looking closer at the FP and FN variants shows that Clair is approaching the limit on how accurately it can call variants using pileup data. Some of the erroneous variant calls can be corrected using complete read alignments instead of pileup data. However, dealing with complete read alignments requires a more powerful neural network design with much greater computational demands. In the future, we will explore using an ensemble method to handle the majority of the variants using Clair, while for the extremely tricky ones we will use a new, more sophisticated method.

The quality and sufficiency of training data is key to the performance of Clair, as well as other deep learning based variant callers, such as DeepVariant. To train a model for production purposes, we used five samples (HG001 to 5) for Illumina data, but only two samples (HG001 and HG002) for ONT, due to the limited availability of public high-coverage whole genome sequencing datasets for the GIAB samples. ONT sequencing of the other GIAB samples is ongoing, and more data will be available in the near future. With additional datasets, we expect to see even higher performance in Clair on ONT data.

On ONT data, although Clair performed the best, its indel calling precision and recall were only about 80%, even excluding GA4GH low-complexity regions, which leaves substantial room for improvement. While the precision can be further improved by considering complete read alignments, the recall is bounded by input and can be improved only with a lower read-level base-calling error rate. Future improvements in ONT technology offer the possibility of reducing the error rate to 2-3%, which in turn should improve Clair’s ability to detect indels in these data.

The GIAB datasets we used for model training have moderate whole-genome sequencing coverage. Although we can use samples with very high coverage (over 300-fold, which is sometimes seen in amplicon sequenced data) with Clair for variant calling, such samples might show degraded performance because very high coverage variants were not adequately observed in model training. To solve this problem, we propose two methods. One method is to do transfer learning using a trained model on additional datasets with very high coverage. Clair supports transfer learning and can be applied to additional datasets instantly. Another method is an ensemble method, which generates multiple copies of randomly subsampled read alignments at a candidate variant for Clair to call variant. A majority vote or a decision tree can be used to make the final decision, using the results of each copy.

A limitation of Clair is that it cannot be applied to polyploid species, which are inconsistent with its neural network design. For the same reasons, Clair is not applicable to somatic variant calling, where a single sample might hold multiple distinct populations of cells. Our next steps include extending Clair to support polyploid species and somatic variant calling.

## Method

### Clair’s input/output

#### Input

For a truth variant for training or a candidate variant for calling, the read alignments that overlap or are adjacent to the variant are summarized (i.e. pile-up data) into a three-dimensional tensor of shape 33 by 8 by 4, comprising 1056 integer numbers. The three dimensions correspond to the position, the count of four possible bases from two different strands, and four different ways of counting. In the first dimension, 33 positions include the starting position of a variant at the center and 16 flanking bases on both sides. The second dimension corresponds to the count of ‘A+’, ‘A-’, ‘C+’, ‘C-’, ‘G+’, ‘G-’, ‘T+’ or ‘T-’, with the symbols +/-denoting the count from the forward/reverse strand. The third dimension replicates the first two dimensions with four different ways of counting to highlight 1) the allelic count of the reference allele, 2) insertions, 3) deletions and 4) single nucleotide alternative alleles. “Supplementary Note – Pseudocode for generating the input tensor” shows the pseudo code of the exact algorithm of how the input tensor is generated. Supplementary Figure 1 demonstrates how the tensors are look like for ONT data at a random ‘non-variant’, a ‘SNP’, an ‘Insertion’, and a ‘Deletion’.

#### Output

The output of Clair has four tasks (a.k.a. four output components, in total 90 probabilities), including 1) the 21-genotype probabilistic model (21 probabilities); 2) zygosity (3 probabilities); 3) the length of the first indel allele (33 probabilities); and 4) the length of the second indel allele (33 probabilities). One of the breakthroughs in Clair is the invention of the 21-genotype probabilistic model. It comprises all of the possible genotypes of a diploid sample at a genome position, including ‘AA’, ‘AC’, ‘AG’, ‘AT’, ‘CC’, ‘CG’, ‘CT’, ‘GG’, ‘GT’, ‘TT’, ‘AI’, ‘CI’, ‘GI’, ‘TI’, ‘AD’, ‘CD’, ‘GD’, ‘TD’, ‘II’, ‘DD’, and ‘ID’, where ‘A’, ‘C’, ‘G’, ‘T’, ‘I’ (insertion) and ‘D’ (deletion) denote the six possible alleles. The new model covers variants with two alternative alleles, which could not be called in Clairvoyante. The zygosity task outputs the probability of the input being 1) a homozygous reference (0/0); 2) heterozygous with 1 or 2 alternative alleles (0/1 or 1/2); or 3) a homozygous variant (1/1). The zygosity task is partially redundant to the 21-genotype task, but it makes decisions independently, and it crosschecks the decision made by the 21-genotype task. Tasks three and four have the same design. They output the length of up to two indel alleles. Each task outputs 33 probabilities, including the likelihood of 1) more than 15bp deleted (<-15bp); 2) any number between −15bp and 15bp, including 0bp, and; 3) more than 15bp inserted (>15bp). In training, the indel allele with a smaller number is set as the first indel allele. For example, for a heterozygous 1bp deletion, the first indel allele is set as −1bp, the second as 0bp (−1bp/0bp). For a heterozygous 1bp insertion, 0bp/1bp is set. This design makes the non-0bp training variants for both tasks balanced. For a heterozygous indel with two alternative alleles, say, one −2bp and one 5bp, −2bp/5bp are set. For a homozygous indel, two indel alleles are set to the same value. For indels longer than 15bp, the exact length is determined using an additional step (Supplementary Note – New methods used in Clair – Dealing with indels longer than 15bp). The output of the two indel allele tasks are also used for crosschecking with the 21-genotype task, with 0bp supporting an SNP allele, and non-0bp supporting an indel allele. More details about how the four tasks crosscheck each other to come up with a result coherently are in “Method – New methods used in Clair – Determining the most probable variant type using the four tasks of Clair”.

### New methods used in Clair

Clair has been fully revamped while a few basic deep-learning techniques in Clairvoyante have been retained, including 1) model initialization; 2) activation function; 3) optimizer; 4) dropout; 5) regularization; and 6) combining multiple samples for model training. Below we discussed the new methods we have applied in Clair.

#### Dealing with indels longer than 15bp

For each candidate variant, Clair directly outputs the length of up to two alternative indel alleles. However, if an insertion goes beyond 15bp, or a deletion goes below −15bp, Clair runs an additional step to decide its exact length and allele. In the additional step, Clair gathers all possible insertion/deletion alleles longer than 15bp at a genome position through pysam (a wrapper around htslib and the samtools^21^ package). Depending on the genotype concluded by Clair, we choose 1) the insertion/deletion with the highest allelic count for ‘AI’, ‘CI’, ‘GI’, ‘TI’, ‘AD’, ‘CD’, ‘GD’ and ‘TD’; 2) the insertions with the highest and/or the second-highest allelic count for ‘II’; 3) the deletions with highest and/or the second-highest allelic count for ‘DD’, or; 4) both the insertion and deletion with the highest allelic count for ‘ID’. The additional step is slow, but it is required only for indels longer than 15bp. We investigated HG001 and found 570,367 indels in its truth variant set; only 10,672 (1.87%) were >15bp. In our experiments, we found the slowdown was acceptable. Users can set an option in Clair to enable this additional step for all indels, but our experiments found that while the improvement in precision is small, it slows down Clair by about two times with Illumina and PacBio CCS data, and by more than 10 times on ONT data.

#### Determining the most probable variant type using the four Clair tasks

Clair outputs data on four tasks. With an independent penultimate layer (Figure 1, FC5 layer) immediately before each task, the output of each task is considered independent. We made two observations from our experiments: 1) for true positive variants, a random task or two will make a mistake occasionally, but usually, the best and the second-best probabilities are near and can be disambiguated if considered with other tasks; 2) for false positive variants, the tasks do not usually agree well with each other, leading to two or more possible decisions with similar probabilities. Thus, in Clair, we implemented a method as a submodule for making a decision using the output of all four tasks. Variants are divided into 10 categories: 1) a homozygous reference allele; 2) a homozygous 1 SNP allele; 3) a heterozygous 1 SNP allele, or heterozygous 2 SNP alleles; 4) a homozygous 1 insertion allele; 5) a heterozygous 1 insertion allele, or heterozygous 1 SNP and 1 insertion alleles; 6) heterozygous 2 insertion alleles; 7) a homozygous 1 deletion allele; 8) a heterozygous 1 deletion allele, or heterozygous 1 SNP and 1 deletion alleles; 9) heterozygous 2 deletion alleles; and 10) a heterozygous 1 insertion and 1 deletion alleles. The likelihood value of the 10 categories is calculated for each candidate variant, and the category with the largest likelihood value is chosen (Pseudocode in “Supplementary Note – Pseudo code for determining the most probable variant type”). The variant quality is calculated as the square of the Phred score of the distance between the largest and the second-largest likelihood values.

#### Cyclical learning rate

The “initial learning rate” and “how the learning rate decays” are two critical hyperparameters in training a deep neural network model. A model might be stuck at a local optimum (i.e. unable to achieve the best precision and recall) if the initial learning rate is too large, or the decay is too fast. But a large initial learning rate, and a slow decay rate make the training process either unstable or take too long to finish. So in common practice, a tediously long grid search that is very costly is needed to find the best hyperparameters. Furthermore, through a grid search, we found that different sequencing technologies differ in their best hyperparameters. This problem makes model training too complicated and largely impedes Clair from being applied to new datasets and sequencing technologies. To solve the problem, we implemented Cyclical Learning Rate (CLR)^22^ in Clair. CLR is a new deep learning technique that eliminates the need to find the best values of the two hyperparameters. CLR gives a way to schedule the learning rate in an efficient way during training, by cyclically varying between a lower and higher threshold. Following the CLR paper, we determined the higher threshold to be 0.03 and the lower threshold to be 0.0001. The two thresholds worked well on the training variants of all three sequencing technologies (Illumina, PacBio CCS and ONT). In terms of which CLR scheduler to use, we chose the triangular schedule with exponential decay. In our experiments, on PacBio CCS and Illumina datasets, CLR decreased model training time by about 1–3 times, while often outperforming the three-step decay method introduced in Clairvoyante for both precision and recall. However, on ONT datasets, CLR has a lower, but almost negligible, performance than the three-step decay. We provide both CLR and three-step decay options in Clair. To train a model for production, we suggest users try both options and choose the best through benchmarking. In our results, we used CLR for PacBio CCS and Illumina datasets, and the three-step decay method for ONT datasets.

#### Focal loss

Our training data uses the truth variants from the GIAB consortium and is unbalanced in terms of variant type. For example, the number of heterozygous variants is nearly twice that of the homozygous variants. SNPs are about five times more numerous than indels. Worst of all, only ∼1.1% (39,898 of 3,619,471 in HG001) of variants have two or more alternative alleles. And among them, only 884 (∼0.024%) are multiallelic SNPs. This problem leads to degenerate models, as the numerous easy variants contribute no useful learning signals and overwhelm training. In our practice, if we leave the problem unaddressed, we observe a significant drop in recall for the underrepresented variant types. For multiallelic SNPs, the recall dropped to zero. To solve this problem, we used the “Focal loss” technique^23^, which applies a modulating term to the cross-entropy loss in Clair’s output to focus training on underrepresented hard variants and down-weight the numerous easy variants. Focal loss calculates the loss as (1 − *p*_*t*_)^*γ*^ × *α*_*t*_ × −log (*p*_*t*_), where *p*_*t*_ = *p, α*_*t*_ = *α*, if the prediction matches the truth, or *p*_*t*_ = (1 − *p*), *α*_*t*_ = (1 − *α*) otherwise. In addition to the traditional cross entropy loss, focal loss uses two more parameters: *γ* (the focusing parameter) to differentiate easy/hard training examples, and *α* (the balancing parameter) to balance the importance of positive/negative training examples. We determined *γ* = 2 and *α* = 0.25 work best for the GIAB truth variants with a 1:2 ratio of truth variant and non-variant. The use of focal loss significantly increases the performance of underrepresented variant types. It also allows us to be more lenient on variant type balance when augmenting the training data.

#### Training data augmentation using subsampled coverage

Lower coverage usually leads to lower precision and recall in variant calling. To train Clair to achieve better performance on variants with lower coverages, we subsampled each dataset into four or nine additional datasets with lower coverages. The subsampling factors *f* are determined as 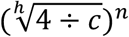, where *c* is full coverage of each sample, 4 is the minimal coverage, h is either 4 or 9, and *n* is from 1 to *h*. Using HG002 as an example, its full coverage is 63.68-fold, and the nine subsampled coverages are 46.82-, 34.43-, 25.31-,18.61-, 13.69-, 10.06-, 7.40-, 5.44- and 4.00-fold. If variant samples were lower than 4x after subsampling, we removed them from training. We used the command “samtools view -s *f*” to generate a subsampled BAM. A different seed counting from zero for random number generation was set for each coverage. The use of subsampled coverages improved the recall on indel significantly.

### Methods tested but showed no improvement to accuracy

In this section we discuss methods we tested that had no effect on Clair’s performance. For researchers working on further improving the performance of Clair, these methods could be avoided or revised.

#### Extend input tensor from 33bp to 49bp and 65bp

Intuitively, a larger input tensor with more flanking bases provides additional information on the surrounding read alignments, which might lead to better precision and recall. Our experiments show that extending the input tensor from 33bp (16bp flanking bases) to 49bp (24bp flanking bases) and 65bp (32bp flanking bases) slows down Clair by 5.4% and 12.6%, respectively. But the improvement was negligible in terms of precision or recall with both SNP and indel.

#### Using non-variants adjacent to true variants as negative samples for model training

Clair, by default, uses a ratio of 1:2 on true variants and non-variants for model training, and the non-variants are randomly selected from the genome, except for the positions with a true variant or insufficient coverage. We experimented using non-variants adjacent to true variants (we tried ±2bp, ±8bp and ±16bp) as negative samples for model training and adjusted the ratio to 1:1:1 on true variants adjacent non-variants and random non-variants. We used adjacent non-variants for training because their input is true variant alike, but a few bases shifted. The hypothesis was that using them as adversarial training samples against the true variants might improve Clair’s performance at high density variants and alignment errors. However, our experiments show that the method decreased recall slightly on both SNP and indel.

#### Incorporating less confident GIAB variants for model training

The GIAB HG001 truth variant dataset includes 3,619,471 truth variants passing all criteria (with the ‘PASS’ tag), and 2,264,796 variants failing one or more criteria. The criteria details were explained by Zook et al. in 2019^13^. Among the failed variants, 310,113 had the ‘allfilteredbutagree’ tag, which means at the same position, the variants called in all the supporting datasets agreed with each other, even though none of them were in the callable regions, in which a range of coverage and minimum alignment quality are met. These variants are considered less confident than those passing all criteria, but might still contribute to training a better model because while a deep neural network can tolerate moderate errors in training data, if any new patterns are provided in additional data, it will be learned by the model and, in turn, improve the performance. We experimented adding the variants with the ‘allfilteredbutagree’ tag to training. However, our results show that the recall went down significantly on SNP, and the precision went down significantly on indel.

#### Discarding homopolymer variants in model training

Variant calling in homopolymer sequences is usually more challenging, and the problem is even worse in SMS technologies since the length of homopolymers is usually underestimated. At longer homopolymers, the signals are usually too discordant, so it is common for humans to make mistakes with them. From the feature engineering point of view, variants in homopolymer sequences are confusing and less informative, and might lead to a degenerate model. We tested model training without variants at homopolymer sequences longer than 5bp. Our results show that both precision and recall degrade significantly if homopolymer variants are not used in model training.

### Benchmarking

#### The GIAB truth variant datasets

We used the GIAB version 3.3.2 datasets as our truth variants. Depending on the availability of deep sequencing data, our ONT experiments used samples HG001 or HG001+HG002 for model training, our PacBio CCS experiments used HG001 or HG001+HG005, and our Illumina experiments used HG001 or HG001+HG003+HG004+HG005. For benchmarking, ONT, PacBio CCS and Illumina experiments have used HG002, HG005, and HG002, respectively. The links to the truth variants and high-confidence regions are available in “Methods – Data sources – Truth variants”. Depending on the reference genome used in the already available read alignments, we used GRCh38 for our ONT and Illumina experiments, and GRCh37 for our PacBio CCS experiments. The links to the reference genomes we used are available in “Methods – Data sources – Reference genomes”

#### Removing GA4GH low-complexity regions from benchmarking

Krusche et al.^6^ from the GA4GH benchmarking team and the GIAB consortium published the low-complexity regions, including homopolymers, STRs, VNTRs, and other repetitive sequences for stratifying variants in their paper titled “Best practices for benchmarking germline small-variant calls in human genomes”. In the low-complexity regions larger than 10bp, ONT’s performance degraded significantly (precision −11.41%, recall −55.33%), while that of PacBio CCS and Illumina dropped only 0.99–1.67% in precision and recall (Supplementary Table 5). Thus, when computing variant calling using ONT, we suggest removing the variants called in the low-complexity regions. In our benchmarks for all datasets, in addition to using the high-confidence regions of each sample provided by GIAB, we removed the low-complexity regions. The procedures are available in “Supplementary Note – Commands – Remove GA4GH low complexity regions from GIAB’s high-confidence regions”. There was retention of 92.61–93.47% high-confidence regions in GRCh38, and 94.40–95.05% in GRCh37 of the five samples HG001 to 5 after removing the low-complexity regions (Supplementary Table 8).

#### Benchmarking methods and metrics

Clair trains a model either for 30 epochs, using the Cyclical Learning Rate (used for PacBio CCS and Illumina datasets), or by decaying the learning rate three times (by one tenth each time) until the validation losses converge (used for ONT datasets). While the performance of last few epochs are generally similar, the best-performing one will be chosen for benchmarking. We did not run replications of model training because choosing from the best epoch actually resembles the process of having multiple replications. In ONT and Illumina experiments, the GRCh38 reference genome was used, while in PacBio CCS experiments, GRCh37 was used. For each variant calling experiment, we used the submodule vcfeval in RTG Tools^24^ version 3.9 to generate three metrics, ‘Precision’, ‘Recall’, and ‘F1-score’, for five categories of variants: ‘Overall’, ‘SNP’, ‘Indel’, ‘Insertion’, and ‘Deletion’. All time consumptions were gauged on two 12-core Intel Xeon Silver 4116 (in total 24 cores), with 12 concurrent Clair processes, each with 4 Tensorflow threads. As Clair has some serial steps that use only one thread, we observed our setting sufficient to maximize the utilization of all 24 cores. For other variant callers, including DeepVariant, Longshot and Medaka, options were to set to use all 24 cores for the best speed.

### Computational performance

Clair requires Python3, Pypy3 and Tensorflow. Variant calling using Clair requires only a CPU. For a typical 30-fold human WGS sample, Clair takes about an hour for Illumina data and PacBio CCS data, and five hours on ONT data, using two 12-core Intel Xeon Silver 4116 processors. Memory consumption depends on both input data and concurrency. ONT data has a higher memory footprint than Illumina and PacBio CSS, while Clair is capped at 7GB per process (helper scripts at 4.5GB and Tensorflow at 2.5GB). Model training requires a high-end GPU; we used the Nvidia Titan RTX 24GB in our experiment. Using Clair’s default parameters, generating 1 million training samples takes about 38 seconds. For example, the Illumina model with four samples (HG001, 3, 4, 5) and 30 coverages in total (10 for 1 and 5, 5 for 2 and 3) has 284,367,735 training samples and takes about 11,000 seconds per epoch. In comparison, the Nvidia RTX 2080 Ti 11GB is about 15% slower, and the Nvidia GTX 1080 Ti 11GB is about 35% slower.

## Code availability

Clair is open source, available at https://github.com/HKU-BAL/Clair.

## Data availability

The authors declare that all data supporting the findings of this study are available at the links in the paper and its supplementary information files.

## Acknowledgements

We thank Steven Salzberg, Mike Schatz, and Fritz Sedlazeck for their valuable comments. R. L. was supported by the ECS (Grant No. 27204518) of the HKSAR government, and the URC fund at HKU. T. L., C. W., Y. W., C. T., C. Li. and C. Le. were supported by the ITF (Grant No. ITF/331/17FP) from the Innovation and Technology Commission, HKSAR government.

## Author contributions

R. L. and T. L. conceived the study. R. L, C. W., Y. W., C. T., C. Li. and C. Le. analyzed the data and wrote the paper.

## References

1 Goodwin, S., McPherson, J. D. & McCombie, W. R. Coming of age: ten years of next-generation sequencing technologies. Nat Rev Genet 17, 333–351, doi: 10.1038/nrg.2016.49 (2016).

2 Ashley, E. A. Towards precision medicine. Nat Rev Genet 17, 507–522, doi: 10.1038/nrg.2016.86 (2016).

3 Li, H. Toward better understanding of artifacts in variant calling from high-coverage samples. Bioinformatics 30, 2843–2851, doi: 10.1093/bioinformatics/btu356 (2014).

4 Luo, R., Schatz, M. C. & Salzberg, S. L. 16GT: a fast and sensitive variant caller using a 16-genotype probabilistic model. GigaScience (2017).

5 Van der Auwera, G. A. et al. From FastQ data to high confidence variant calls: the Genome Analysis Toolkit best practices pipeline. Curr Protoc Bioinformatics 43, 11 10 11–33, doi: 10.1002/0471250953.bi1110s43 (2013).

6 Krusche, P. et al. Best practices for benchmarking germline small-variant calls in human genomes. Nat Biotechnol 37, 555–560, doi: 10.1038/s41587-019-0054-x (2019).

7 The long view on sequencing. Nat Biotechnol 36, 287, doi: 10.1038/nbt.4125 (2018).

8 Sedlazeck, F. J., Lee, H., Darby, C. A. & Schatz, M. C. Piercing the dark matter: bioinformatics of long-range sequencing and mapping. Nat Rev Genet, doi: 10.1038/s41576-018-0003-4 (2018).

9 Ameur, A., Kloosterman, W. P. & Hestand, M. S. Single-Molecule Sequencing: Towards Clinical Applications. Trends Biotechnol 37, 72–85, doi: 10.1016/j.tibtech.2018.07.013 (2019).

10 Luo, R., Sedlazeck, F. J., Lam, T. W. & Schatz, M. C. A multi-task convolutional deep neural network for variant calling in single molecule sequencing. Nat Commun 10, 998, doi: 10.1038/s41467-019-09025-z (2019).

11 Zook, J. M. et al. Extensive sequencing of seven human genomes to characterize benchmark reference materials. Sci Data 3, 160025, doi: 10.1038/sdata.2016.25 (2016).

12 Zook, J. M. et al. Integrating human sequence data sets provides a resource of benchmark SNP and indel genotype calls. Nat Biotechnol 32, 246–251, doi: 10.1038/nbt.2835 (2014).

13 Zook, J. M. et al. An open resource for accurately benchmarking small variant and reference calls. Nature Biotechnology 37, 561–566, doi: 10.1038/s41587-019-0074-6 (2019).

14 Jain, M. et al. Nanopore sequencing and assembly of a human genome with ultra-long reads. Nat Biotechnol 36, 338–345, doi: 10.1038/nbt.4060 (2018).

15 Edge, P. & Bansal, V. Longshot enables accurate variant calling in diploid genomes from single-molecule long read sequencing. Nat Commun 10, 4660, doi: 10.1038/s41467-019-12493-y (2019).

16 medaka: Sequence correction provided by ONT Research. https://github.com/nanoporetech/medaka, accessed Nov 17 2019.

17 Wenger, A. M. et al. Accurate circular consensus long-read sequencing improves variant detection and assembly of a human genome. Nat Biotechnol 37, 1155–1162, doi: 10.1038/s41587-019-0217-9 (2019).

18 Poplin, R. et al. A universal SNP and small-indel variant caller using deep neural networks. Nature Biotechnology 2018/09/24/online, doi: 10.1038/nbt.4235 (2018).

19 Simpson, J. T. et al. Detecting DNA cytosine methylation using nanopore sequencing. Nature methods 14, 407 (2017).

20 Poplin, R. et al. DeepVariant training data. https://github.com/google/deepvariant/blob/r0.9/docs/deepvariant-details-training-data.md, accessed Nov 22 2019.

21 Li, H. et al. The Sequence Alignment/Map format and SAMtools. Bioinformatics 25, 2078–2079, doi: 10.1093/bioinformatics/btp352 (2009).

22 Smith, L. N. in 2017 IEEE Winter Conference on Applications of Computer Vision (WACV). 464–472 (IEEE).

23 Lin, T.-Y., Goyal, P., Girshick, R., He, K. & Dollár, P. in Proceedings of the IEEE international conference on computer vision. 2980–2988.

24 Cleary, J. G. et al. Joint variant and de novo mutation identification on pedigrees from high-throughput sequencing data. J Comput Biol 21, 405–419, doi: 10.1089/cmb.2014.0029 (2014).

