## Supplementary Materials for "Clair: Exploring the limit of using a deep neural network on pileup data for germline variant calling"

Ruibang Luo, Chak-Lim Wong, Yat-Sing Wong, Chi-Ian Tang, Chi-Man Liu, Chi-Ming Leung, Tak-Wah Lam

### Supplementary Notes

#### Pseudocode for generating the input tensor

```
Input: p, genome position of a candidate variant, 1-based
       r, reference genome
       a, a set of elements representing the read alignments covering p at bp level,
          each element contains four members:
          1) the matched reference position (refpos), 1-based,
          2) the offset of this bp within its insertion pattern (insoffset),
             1-based (only applicable if this bp is an inserted base),
          3) the bp (IUPAC nucleotide A/C/G/T/U/R/Y/S/W/K/M/B/D/H/V) or the gap '-' in r at refpos (refbase),
          4) the bp or the gap in the corresponding aligned read at readpos (readbase).

Output: A matrix x of order 33 by 8 by 4

h.insert('A', 0)
h.insert('C', 1)
h.insert('G', 2)
h.insert('T', 3)
h.insert('U', 3)
h.insert('R', 0)
h.insert('Y', 1)
h.insert('S', 1)
h.insert('W', 0)
h.insert('K', 2)
h.insert('M', 0)
h.insert('B', 1)
h.insert('D', 0)
h.insert('H', 0)
h.insert('V', 0)

create a zero-filled matrix x of size (33 x 8 x 4)

for each item in a do:
    /* Skip the Ns in reference */
    if a[item].refbase not one of "ACGTURYSWKMBDHV-" then:
        continue

    /* Skip the Ns in read */
    if a[item].readbase not one of "ACGTURYSWKMBDHV-" then:
        continue

    /* Center the candidate variant in the first dimension (33) of x */
    offset <- refpos - p + 16

    /* Skip if beyond scope */
    if offset < 0 or offset > 32 then:
        continue

    /* Translate refbase and readbase into index */
    refbaseIndex <- h.get(a[item].refbase)
    readbaseIndex <- h.get(a[item].readbase)

    /* Calculate strand offset, positive +0, negative +4*/
    strandOffset <- 0
    if a[item].isNegativeStrand then:
        strandOffset <- 4

    /* If not a deleted base */
    if a[item].readbase ≠ '-' then:
        /* If not an inserted base either, i.e. a ref or a SNP */
        if a[item].refbase ≠ '-' then:
            increase x[offset][refbaseIndex + strandOffset][0] by 1
            increase x[offset][readbaseIndex + strandOffset][1] by 1
            increase x[offset][refbaseIndex + strandOffset][2] by 1
            increase x[offset][readbaseIndex + strandOffset][3] by 1
        /* If is an inserted base */
        if a[item].refbase = '-' then:
            newOffset <- min(offset + a[item].insoffset, 32)
            increase x[newOffset][readbaseIndex + strandOffset][1] by 1
    /* If is a deleted base */
    if a[item].readbase = '-' then:
        increase x[offset][refbaseIndex + strandOffset][2] by 1

for i = 0 to 32 do:
    for j = 0 to 7 do:
        x[i][j][1] <- x[i][j][1] - x[i][j][0]
        x[i][j][2] <- x[i][j][2] - x[i][j][0]
        x[i][j][3] <- x[i][j][3] - x[i][j][0]

return x
```

#### Pseudocode for determining the most probable variant type

This section describes how to determine the variant to be reported (including the indel sequences, if any) given the output tensors of Clair’s deep network.

**Input**

(1) The reference allele *r*
(2) Network output probabilities:
    (a) gt21: bi-allelic GT21 probabilities (21 values)
    (b) zygosity: “ref / hom / het” for the probabilities of homozygous reference, homozygous variant and heterozygous variant respectively (3 values)
    (c) indel_length1: first length of indel, between -16 and +16 (33 values)
    (d) indel_length2: second length of indel, between -16 and +16 (33 values)

**Compute likelihood variables**

For each likelihood variable, compute its value by multiplying the corresponding output probabilities according to the following table. For example,

HomDel(8) = gt21[DD] × zygosity[hom] × indel_length1[-8] × indel_length2[-8]

| Type | Likelihood variable | gt21 | zygo-  sity | indel_  length1,2 | Remark |
| --- | --- | --- | --- | --- | --- |
| Ref | HomRef | *rr* | ref | (0, 0) | reference allele *r* |
| SNP | HomSNP(*x*) | *xx* | hom | (0, 0) | *x* ∈ {A,C,G,T}, *x* ≠ *r* |
|  | HetSNP(*x*, *y*) | *xy* | het | (0, 0) | *x*, *y* ∈ {A,C,G,T}, *x* ≠ *y*, *x* ≠ *r*, *y* ≠ *r* |
| Ins | HomIns(*L*) | II | hom | (*L*, *L*) | 1 ≤ *L* ≤ 16 |
|  | HetOneIns(*x*, *L*) | *x*I | het | (0, *L*) * | *x* ∈ {A,C,G,T}; 1 ≤ *L* ≤ 16 |
|  | HetTwoIns(*L*_1_, *L*_2_) | II | het | (*L*_1_, *L*_2_) * | 1 ≤ *L*_1_ ≤ *L*_2_ ≤ 16 |
| Del | HomDel(*L*) | DD | hom | (-*L*, -*L*) | 1 ≤ *L* ≤ 16 |
|  | HetOneDel(*x*, *L*) | xD | het | (-*L*, 0) * | *x* ∈ {A,C,G,T}; 1 ≤ *L* ≤ 16 |
|  | HetTwoDel(*L*_1_, *L*_2_) | DD | het | (-*L*_1_, -*L*_2_) * | 1 ≤ *L*_1_ < *L*_2_ ≤ 16 |
| InsDel | HetInsDel(*L*_1_, *L*_2_) | ID | het | (-*L*_1_, *L*_2_) * | 1 ≤ *L*_1_, *L*_2_ ≤ 16 |

(*) For variables with two different indel lengths *M* ≠ *N*, we choose the order among (*M*, *N*) and (*N*, *M*) which gives the larger value.

**Determine the most probable variant type**

The procedure is as follows.

1. Find the likelihood variable with the largest value.
2. If it is of type Ref, we report no variant.
3. Otherwise, if it is of type SNP, we report the corresponding SNP.
4. Otherwise, it involves indels. We try to extract the insertion or deletion sequence (or two sequences in case it is heterozygous) accordingly:
   a. If the option pysam_for_all_indel_length is **disabled**:
      i. If the specified indel length *L* satisfies -15 ≤ *L* ≤ 15:
         1. If the specified indel type is insertion, we infer the insertion sequence from the input tensor
         2. If the specified indel type is deletion, we infer the deletion sequence from the reference sequence and the indel length
      ii. Otherwise, *L* = ±16, which is a special value for representing indels longer than 15 bp. In this case, we do not know the indel length, therefore we resort to checking the read alignment for the indel sequence:
          1. Find all reads supporting the specified indel type (insertion or deletion) with indel length > 15 bp
          2. If at least one supporting read can be found, we extract the most common indel sequence from all supporting reads; otherwise, we infer the indel sequence from the input tensor as in Step 4(a)(i)
   b. If the option pysam_for_all_indel_length is **enabled**:
      i. Find all reads supporting the specified indel type (insertion or deletion) and indel length
      ii. If at least one supporting read can be found, we extract the most common indel sequence from all supporting reads; otherwise, extraction of the indel sequence fails (we will *not* infer it from the input tensor)
5. If all the required indel sequences are successfully extracted, we report the corresponding variant. Otherwise, we discard this likelihood variable and go back to Step 1.

#### Command

##### Remove GA4GH low complexity regions from GIAB's high-confidence regions

###### GRCh38

Download the following bed files from <https://github.com/jzook/genome-data-integration/tree/master/NISTv3.3.2/filtbeds/GRCh38>

AllRepeats_lt51bp_gt95identity_merged.bed

AllRepeats_51to200bp_gt95identity_merged.bed

AllRepeats_gt200bp_gt95identity_merged.bed

SimpleRepeat_imperfecthomopolgt10_slop5.chrPrefix.bed

remapped_superdupsmerged_all_sort.bed

remapped_PacBio_MetaSV_svclassify_mergedSVs.bed

hg38_self_chain_nosamepos_withalts_gt10k.bed

GCA_000001405.15_GRCh38_no_alt_plus_hs38d1_analysis_set_REF_N.bed

For the GRCh38 high-confidence regions bed file $j in each sample in HG001, 2, 3, 4 and 5, do:

cat $j.bed | \

bedtools subtract -a stdin -b AllRepeats_lt51bp_gt95identity_merged.bed | \

bedtools subtract -a stdin -b AllRepeats_51to200bp_gt95identity_merged.bed | \

bedtools subtract -a stdin -b AllRepeats_gt200bp_gt95identity_merged.bed | \

bedtools subtract -a stdin -b SimpleRepeat_imperfecthomopolgt10_slop5.chrPrefix.bed | \

bedtools subtract -a stdin -b remapped_superdupsmerged_all_sort.bed | \

bedtools subtract -a stdin -b remapped_PacBio_MetaSV_svclassify_mergedSVs.bed | \

bedtools subtract -a stdin -b hg38_self_chain_nosamepos_withalts_gt10k.bed | \

bedtools subtract -a stdin -b GCA_000001405.15_GRCh38_no_alt_plus_hs38d1_analysis_set_REF_N.bed > $j.clean.bed

###### GRCh37

Download the following bed files from <https://github.com/jzook/genome-data-integration/tree/master/NISTv3.3.2/filtbeds/GRCh37>

AllRepeats_lt51bp_gt95identity_merged_slop5.bed

AllRepeats_51to200bp_gt95identity_merged_slop5.bed

AllRepeats_gt200bp_gt95identity_merged_sort.bed

SimpleRepeat_imperfecthomopolgt10_slop5.bed

example_of_no_ref_regions_input_file_b37.bed

superdupsmerged_all_sort.bed

mm-2-merged.bed

For the GRCh35 high-confidence regions bed file $j in each sample in HG001, 2, 3, 4 and 5, do:

cat $j.bed | \

bedtools subtract -a stdin -b AllRepeats_lt51bp_gt95identity_merged_slop5.bed | \

bedtools subtract -a stdin -b AllRepeats_51to200bp_gt95identity_merged_slop5.bed | \

bedtools subtract -a stdin -b AllRepeats_gt200bp_gt95identity_merged_sort.bed | \

bedtools subtract -a stdin -b SimpleRepeat_imperfecthomopolgt10_slop5.bed | \

bedtools subtract -a stdin -b example_of_no_ref_regions_input_file_b37.bed | \

bedtools subtract -a stdin -b superdupsmerged_all_sort.bed | \

bedtools subtract -a stdin -b mm-2-merged.bed > $j.clean.bed

##### Running other variant callers

###### Clairvoyante (v1.02)

###### Variant calling

```
python clairvoyante/callVarBamParallel.py \
       --chkpnt_fn trainedModels/fullv3-pacbio-ngmlr-hg001+hg002+hg003+hg004-hg19 \
       --ref_fn ref.fasta \
       --bam_fn input.bam \
       --sampleName HG005 \
       --output_prefix hg005 \
       --threshold 0.2 \
       --minCoverage 4 \
       --tensorflowThreads 4 > commands.sh
```

*# The commands in the commands.sh script can be run in parallel.*

*# For each experiment, a model was selected according to the sequencing technology. The details of available models are at https://github.com/aquaskyline/Clairvoyante*

###### Model training

*# For commands to train a Clair model, please refer the training section in* [*https://github.com/HKU-BAL/Clair/blob/master/README.md*](https://github.com/HKU-BAL/Clair/blob/master/README.md)

###### DeepVariant (v0.8)

###### Variant calling

```
sudo docker run \
  -v "input_dir":/input \
  -v "output_dir":/output \
  gcr.io/deepvariant-docker/deepvariant:"0.8.0" \
  /opt/deepvariant/bin/run_deepvariant \
  --model_type=WGS \
  --ref=ref.fasta \
  --reads=input.bam \
  --output_vcf=/output/output.vcf.gz \
  --output_gvcf=/output/output.g.vcf.gz \
  --num_shards=24
```

###### Longshot (v0.3.4)

###### Variant calling

seq 22 | parallel "longshot -A -r {} --bam input.bam \

--strand_bias_pvalue_cutoff 0.01 \

--ref ref.fasta \

--out output_{}.vcf"

###### Medaka (v.0.10.0)

###### Remove read group

samtools view -H input.bam | grep -v "^@RG" | samtools reheader - input.bam > no_read_group.bam

###### Index bam

samtools index no_read_group.bam

###### Variant calling

medaka_variant -f ref.fa -i no_read_group.bam

#### Data Sources

##### Reference genomes

###### GRCh38

<ftp://ftp.ncbi.nlm.nih.gov/genomes/all/GCA/000/001/405/GCA_000001405.15_GRCh38/seqs_for_alignment_pipelines.ucsc_ids/GCA_000001405.15_GRCh38_no_alt_plus_hs38d1_analysis_set.fna.gz>

###### GRCh37

<ftp://ftp.1000genomes.ebi.ac.uk/vol1/ftp/technical/reference/phase2_reference_assembly_sequence/hs37d5.fa.gz>

##### Truth Variants (Genome in a Bottle dataset version 3.3.2)

###### HG001 (NA12878), GRCh38

<ftp://ftp-trace.ncbi.nlm.nih.gov/giab/ftp/release/NA12878_HG001/NISTv3.3.2/GRCh38>

###### HG001 (NA12878), GRCh37

<ftp://ftp-trace.ncbi.nlm.nih.gov/giab/ftp/release/NA12878_HG001/NISTv3.3.2/GRCh37>

###### HG002 (NA24385), GRCh38

<ftp://ftp-trace.ncbi.nlm.nih.gov/giab/ftp/release/AshkenazimTrio/HG002_NA24385_son/NISTv3.3.2/GRCh38>

###### HG002 (NA24385), GRCh37

<ftp://ftp-trace.ncbi.nlm.nih.gov/giab/ftp/release/AshkenazimTrio/HG002_NA24385_son/NISTv3.3.2/GRCh37>

###### HG005 (NA24631), GRCh38

<ftp://ftp-trace.ncbi.nlm.nih.gov/giab/ftp/release/ChineseTrio/HG005_NA24631_son/NISTv3.3.2/GRCh38/>

###### HG005 (NA24631), GRCh37

<ftp://ftp-trace.ncbi.nlm.nih.gov/giab/ftp/release/ChineseTrio/HG005_NA24631_son/NISTv3.3.2/GRCh37/>

##### Oxford Nanopore (ONT) Data

###### HG001 (NA12878) rel6, GRCh38, ~44.3-fold

###### Raw data

<http://s3.amazonaws.com/nanopore-human-wgs/rel6/rel_6.fastq.gz>

###### Descriptions on rel6 data

<https://github.com/nanopore-wgs-consortium/NA12878/blob/771e8544dd985a1c98a6256c60aedcc23a5a46c2/Genome.md>

###### Alignments, GRCh38

<http://www.bio8.cs.hku.hk/clairvoyante/orginalData_hg001/ont-hg38-minimap2-rel6/rel6_hs38d1.sorted.bam>

###### HG001 (NA12878) Phillip, ~124.1-fold

*# The raw reads were provided by Philipp Rescheneder through personal communication. The reads were basecalled with Guppy 2.2.3 using the flip-flop model. Reference genome hs38d1 and aligner minimap2 2.17-r943-dirty (Li et al, 2017) were used for generating the alignments. The BAM file will be publicly available eventually but is available upon request at this moment. Please write to Philipp Rescheneder for more details.*

###### HG002 (NA24835), GRCh38, ~63.6793-fold

<ftp://ftp-trace.ncbi.nlm.nih.gov/giab/ftp/data/AshkenazimTrio/HG002_NA24385_son/Ultralong_OxfordNanopore/final/ultra-long-ont_GRCh38_reheader.bam>

##### Pacific Bioscience (PacBio) CCS Data

###### HG001 (NA12878), GRCh37, ~30.1164-fold

<ftp://ftp-trace.ncbi.nlm.nih.gov/giab/ftp/data/NA12878/PacBio_SequelII_CCS_11kb/HG001.SequelII.pbmm2.hs37d5.whatshap.haplotag.RTG.trio.bam>

###### HG002 (NA24385), GRCh37, ~33.2069-fold

<ftp://ftp-trace.ncbi.nlm.nih.gov/giab/ftp/data/AshkenazimTrio/HG002_NA24385_son/PacBio_SequelII_CCS_11kb/HG002.SequelII.pbmm2.hs37d5.whatshap.haplotag.RTG.10x.trio.bam>

###### HG005 (NA24631), GRCh37, ~32.7169-fold

<ftp://ftp-trace.ncbi.nlm.nih.gov/giab/ftp/data/ChineseTrio/HG005_NA24631_son/PacBio_SequelII_CCS_11kb/HG005.SequelII.pbmm2.hs37d5.whatshap.haplotag.10x.bam>

##### Illumina Data

###### HG001 (NA12878), GRCh38, ~305.784-fold

<ftp://ftp-trace.ncbi.nlm.nih.gov/giab/ftp/data/NA12878/NIST_NA12878_HG001_HiSeq_300x/NHGRI_Illumina300X_novoalign_bams/HG001.GRCh38_full_plus_hs38d1_analysis_set_minus_alts.300x.bam>

###### HG002 (NA24385), GRCh38, ~51.504-fold

<ftp://ftp-trace.ncbi.nlm.nih.gov/giab/ftp/data/AshkenazimTrio/HG002_NA24385_son/NIST_HiSeq_HG002_Homogeneity-10953946/NHGRI_Illumina300X_AJtrio_novoalign_bams/HG002.GRCh38.300x.bam>

*# We downsampled HG002 to around 52x using “samtools view -s”, the downsampled BAM is at*

<http://www.bio8.cs.hku.hk/clairvoyante/orginalData_hg002/illumina/HG002.GRCh38.50x.rg.bam>

###### HG003 (NA24149), GRCh38, ~57.2978-fold

<ftp://ftp-trace.ncbi.nlm.nih.gov/giab/ftp/data/AshkenazimTrio/HG003_NA24149_father/NIST_HiSeq_HG003_Homogeneity-12389378/NHGRI_Illumina300X_AJtrio_novoalign_bams/HG003.GRCh38.300x.bam>

*# We downsampled HG003 to around 57x using “samtools view -s”, the downsampled BAM is at*

<http://www.bio8.cs.hku.hk/clairvoyante/orginalData_hg003/illumina/HG003.GRCh38.60x.1.bam>

###### HG004 (NA24143), GRCh38, ~65.6567-fold

<ftp://ftp-trace.ncbi.nlm.nih.gov/giab/ftp/data/AshkenazimTrio/HG004_NA24143_mother/NIST_HiSeq_HG004_Homogeneity-14572558/NHGRI_Illumina300X_AJtrio_novoalign_bams/HG004.GRCh38.300x.bam>

*# We downsampled HG004 to around 66x using “samtools view -s”, the downsampled BAM is at*

<http://www.bio8.cs.hku.hk/clairvoyante/orginalData_hg004/illumina/HG004.GRCh38.60x.1.bam>

###### HG005 (NA24631), GRCh38, ~352.34-fold

<ftp://ftp-trace.ncbi.nlm.nih.gov/giab/ftp/data/ChineseTrio/HG005_NA24631_son/HG005_NA24631_son_HiSeq_300x/NHGRI_Illumina300X_Chinesetrio_novoalign_bams/HG005.GRCh38_full_plus_hs38d1_analysis_set_minus_alts.300x.bam>

### Supplementary Tables

#### Supplementary Table 1. ONT benchmarking results.

The table shows the performance of Clair and other variant callers. For Clair, the datasets used for model training and testing are written as ‘*a*:*b*x’, where *a* denotes the suffix of the GIAB sample ID of the dataset, and *b* denotes the coverage of the dataset. The ‘precision’, ‘recall’, and ‘F1-score’ are written in sequence and slash-separated. ‘h37’ means GRCh37 reference genome was used. ‘single cov.’ means only the full coverage was used for model training (i.e., ten subsampled coverages were NOT used for model training). All Clair experiments were set to 1) require a minimum of 0.2 allele frequency; 2) “focal loss” on; 3) “cyclical learning rate” off, and; 3) “pysam for all indels” off.

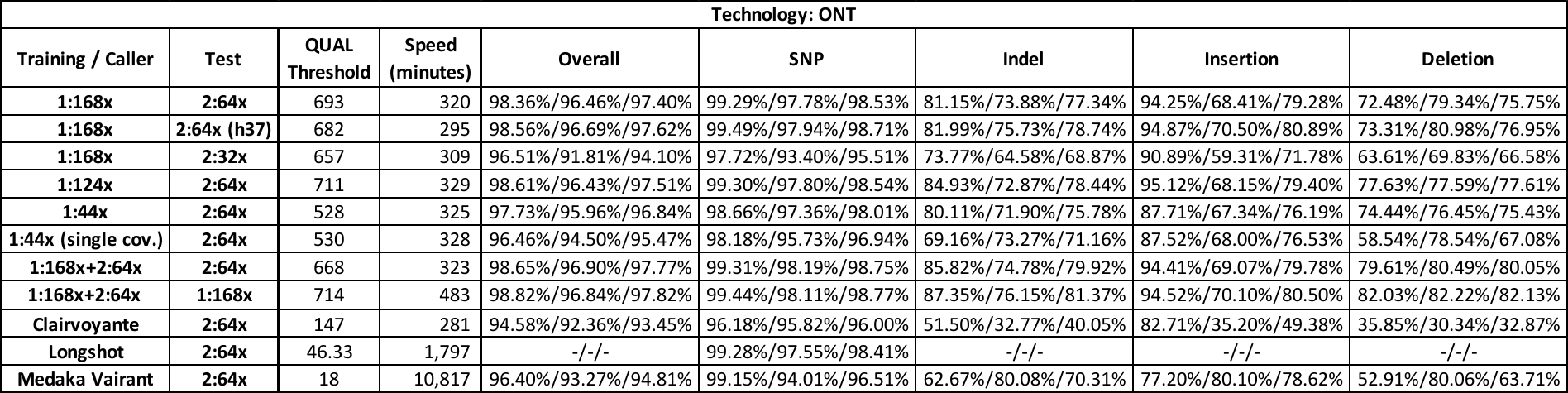

#### Supplementary Table 2. The details of the FP and FN results in ONT experiment 1:168x|2:64x.

In a separate file named "supplementary table 2.xlsx".

#### Supplementary Table 3. PacBio CCS benchmarking results.

The table shows the performance of Clair and other variant callers. For Clair, the datasets used for model training and testing are written as ‘*a*:*b*x’, where *a* denotes the suffix of the GIAB sample ID of the dataset, and *b* denotes the coverage of the dataset. The ‘precision’, ‘recall’, and ‘F1-score’ are written in sequence and slash-separated. All Clair experiments were set to 1) require a minimum of 0.2 allele frequency; 2) “focal loss” on, 3) cyclical learning rate on, and; 4) “pysam for all indels” on.

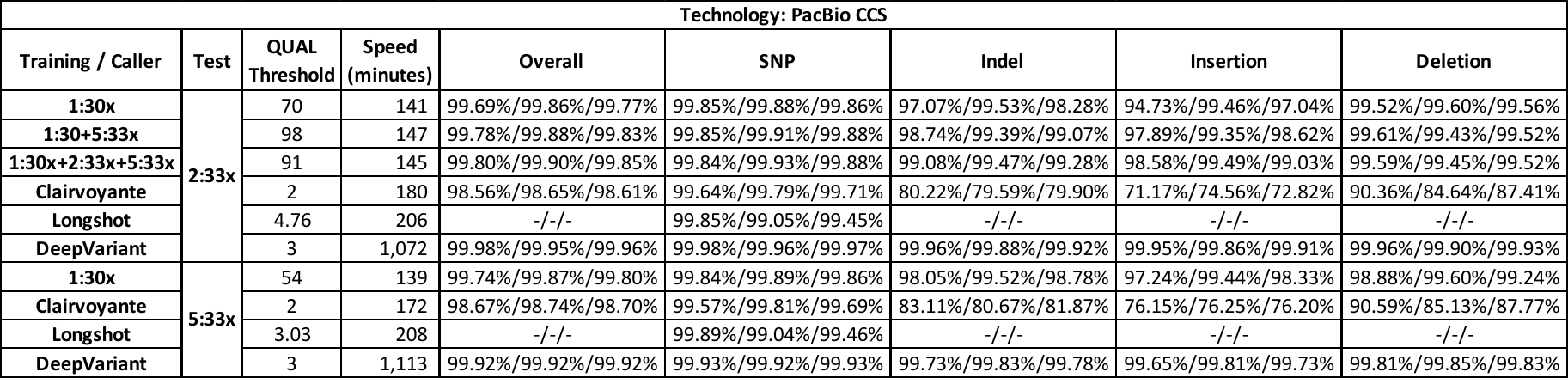

#### Supplementary Table 4. Illumina benchmarking results.

The table shows the performance of Clair and other variant callers. For Clair, the datasets used for model training and testing are written as ‘*a*:*b*x’, where *a* denotes the suffix of the GIAB sample ID of the dataset, and *b* denotes the coverage of the dataset. The ‘precision’, ‘recall’, and ‘F1-score’ are written in sequence and slash-separated. All Clair experiments were set to 1) require a minimum of 0.1 allele frequency; 2) “focal loss” on; 3) “cyclical learning rate” on, and; 3) “pysam for all indels” off.

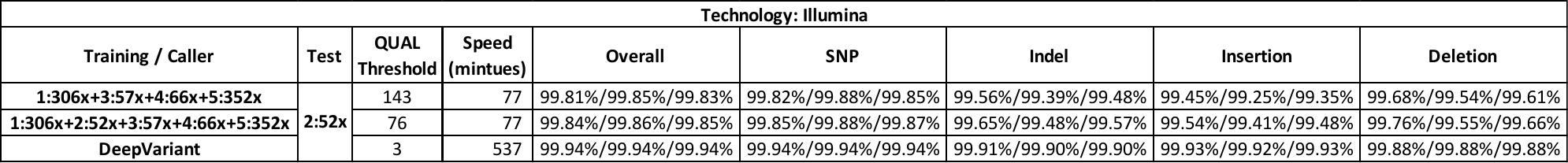

#### Supplementary Table 5. Performance within and without the low complexity regions.

The table shows the performance difference between within and without the GA4GH low complexity regions of different sequencing technologies using Clair. The datasets used for model training and testing are written as ‘*a*:*b*x’, where *a* denotes the suffix of the GIAB sample ID of the dataset, and *b* denotes the coverage of the dataset. The difference in ‘precision’, ‘recall’, and ‘F1-score’ are written in sequence and slash-separated.

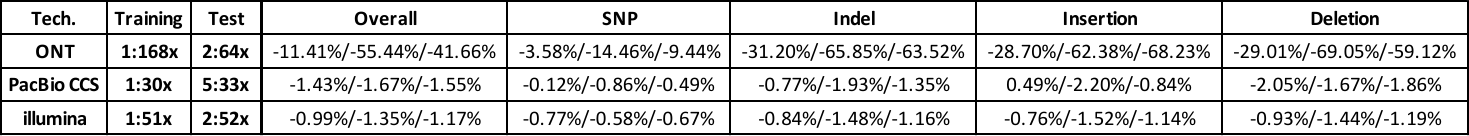

#### Supplementary Table 6. Statistics of removing low complexity regions from benchmarking.

The table shows the number of bases of the 1) high-confidence regions, and; 2) the high-confidence regions less of GA4GH low complexity regions, of five different GIAB samples and two different reference genome versions.

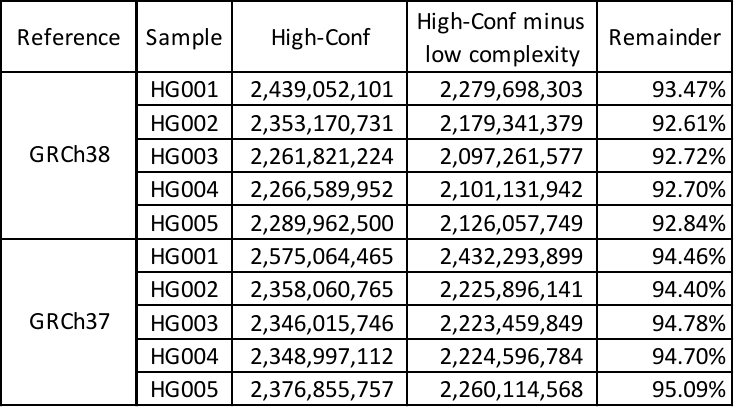

### Supplementary Figures

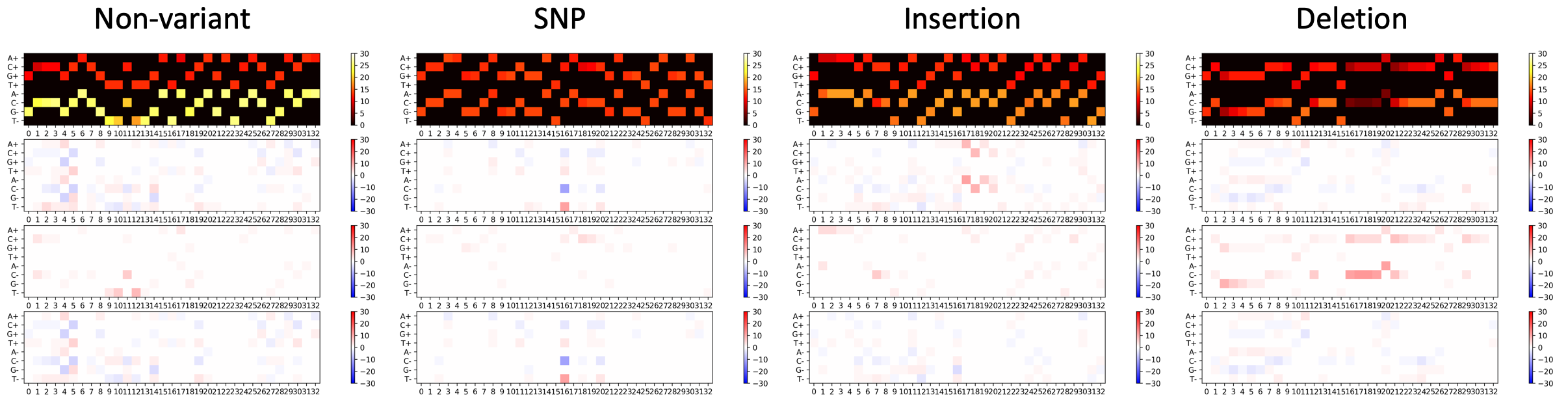

#### Supplementary Figure 1. Demonstration of input tensors on four different variant types

The figure demonstrates how the tensors are look like for ONT data at a random ‘non-variant’, a ‘SNP’, an ‘Insertion’, and a ‘Deletion’. The color intensity represents the strength of a certain signal.
